# Temporo-cerebellar connectivity underlies timing constraints in audition

**DOI:** 10.1101/2021.02.06.430038

**Authors:** Anika Stockert, Michael Schwartze, David Poeppel, Alfred Anwander, Sonja A. Kotz

## Abstract

The flexible and efficient adaptation to dynamic, rapid changes in the auditory environment likely involves generating and updating of internal models. Such models arguably exploit connections between the neocortex and the cerebellum, supporting proactive adaptation. Here we test the functional mechanisms associated with temporo-cerebellar connectivity, verifying these mechanisms for speech sounds. First, we identify lesion-specific deficits for the encoding of short timescale spectro-temporal non-speech and speech properties in patients with left posterior temporal cortex stroke. Second, using lesion-guided probabilistic tractography in healthy participants, we reveal bidirectional temporo-cerebellar connectivity with cerebellar dentate nuclei and crura I/II. These findings imply that the encoding and modeling of rapidly modulated auditory spectro-temporal properties engage a temporo-cerebellar interface. The data further support the conjecture that proactive adaptation to a dynamic environment via internal models is a generalizable principle.

**Significance Statement:** Asymmetric sampling in time, the principle of duration-sensitive hemispheric specialization of the cerebral cortex in the sensory decomposition of sound, is a widely tested hypothesis in auditory neuroscience. This functional organization is mirrored in the cerebellar cortex, implicated in the internal forward-modeling of sensory feedback that arises from motor actions. The potential structural and functional integration of these systems is not well understood. Using a unique combination of causal lesion-symptom mapping in persons with temporal lobe damage and diffusion-weighted magnetic resonance neuroimaging in healthy persons, we identify one key missing link and provide evidence for cross-lateral temporo-cerebellar connectivity using probabilistic white matter fiber tractography. These cerebellar-pontine-temporal cortex connections not only support asymmetric sampling in time but also establish the basis for a generalizable role of cerebellar forward-modeling in sensation beyond the monitoring of sensory feedback in the motor domain.

## Introduction

Current theories of motor control postulate that the cerebellum plays a foundational role in monitoring motor performance and its sensory consequences (Wolpert, Ghahramani, & Jordan, 1995; Wolpert & Miall, 1996). This important concept has been extended to anticipatory sensory and cognitive processes (Ito, 2008; Ramnani, 2006). In this view, cortico-cerebellar interfaces implement essential properties of motor and non-motor (internal) models, i.e., representations that can be used to anticipate future events, thereby maximizing the precision of motor, sensory, and cognitive performance (Ito, 2008). A particularly salient attribute of such models, under active study, concerns timing. The *cerebellar timing hypothesis* claims that the cerebellum encodes the precise temporal locus of sensory events (Ivry, Spencer, Zelaznik, & Diedrichsen, 2002; Spencer & Ivry, 2013), with potential asymmetric hemispheric sensitivities. While the right cerebellar hemisphere prefers rapid, the left prefers slow signal modulations (Callan, Kawato, Parsons, & Turner, 2007).

This cerebellar structural and functional organization mirrors the asymmetric sampling of non-speech (Boemio, Fromm, Braun, & Poeppel, 2005; Zatorre & Belin, 2001) and speech sounds (Poeppel, 2003) in auditory cortex. Concretely, the sampling of continuous sounds has been argued to proceed in time-windows of different lengths, which, in turn, translate into different linguistic segments of speech (e.g., phonemes and syllables) (Flinker, Doyle, Mehta, Devinsky, & Poeppel, 2019). Thus, anticipatory modeling of shorter and longer segments may pave the way for optimal sound and speech perception.

If the cerebellum interacts with areas in the cerebral cortex to implement internal models that reflect the temporal structure of perceptual experience, this asymmetry requires a pattern of cross-lateral right-cerebellar-left-cortical and left-cerebellar-right-cortical connectivity. Interestingly, this neurofunctional connectivity pattern is evident in the *motor* domain. Similar observations in non-motor domains remain largely unexplored. Is the representation and analysis of temporal information a general feature underpinning cortico-cerebellar functional connectivity? The major goal of this suite of experiments is to fill this gap in our understanding by combining functional and structural prior knowledge with new empirical evidence in a multidimensional approach to systematically contrast fast and slow temporal modulations of sound. The study is anchored in lesion data, which establish the most direct link between function and structure.

Existing deficit-lesion data show that damage to the cerebellum impairs the perception of temporal voicing contrasts (Ackermann, Graber, Hertrich, & Daum, 1997), duration judgments of intervals (Ivry & Keele, 1989), and the ability to use temporal event structure to update a representation of the auditory environment (Kotz, Stockert, & Schwartze, 2014). Similarly, left temporal cortex lesions, in particular, lead to impairments of temporal order judgments (Efron, 1963; Swisher & Hirsh, 1972), discrimination of rapidly presented complex tone pairs (micropatterns; Chedru, Bastard, and Efron, 1978), phonological discrimination associated with increased detection thresholds for rapid (but not slow) modulations (Robson, Grube, Lambon Ralph, Griffiths, & Sage, 2013), detection of short time-scale voicing contrasts, and increased temporal order thresholds (Fink, Churan, & Wittmann, 2006). Together these findings confirm the functional relevance of differential temporal sensitivities in *both* cerebellar and temporal cortex. This evidence then motivates the question of how cerebellum and temporal cortex interface to optimize the processing of spectro-temporal information at different timescales in audition (Boemio et al., 2005; Kotz & Schwartze, 2010; Poeppel, 2003).

Here we combine lesion mapping, tractography, and behavioral data to gain new mechanistic insight. First, patients, who suffered from a circumscribed stroke in the left posterior superior temporal sulcus - with spared Heschl’s gyrus, i.e., putative primary auditory cortex – are characterized. Based on an extensive literature, such patients are expected to show impaired temporal discrimination for non-verbal and verbal information, restricted to fast modulations, such as voicing and place of articulation contrasts (Boemio et al., 2005; Elangovan & Stuart, 2008; Rosen, 1992). We test this hypothesis using a range of speech and non-speech materials. Second, we predict that specific lesion-symptom mapping will identify a critical seed region to distinguish the well documented dorsal and ventral fiber tracts of the temporo-frontal speech network (Frey, Campbell, Pike, & Petrides, 2008; Friederici, 2011; Saur et al., 2008; Turken & Dronkers, 2011). Third, and most critically, if the generalized timing conjecture is on the right track, anatomic tractrography should reveal direct contralateral connections linking the left posterior temporal cortex with the right posterior lateral cerebellum (cerebellar crura I/II), ostensibly engaged in auditory processing (Petacchi, Laird, Fox, & Bower, 2005).

Modeling and adapting to a dynamic auditory environment require a sufficiently detailed representation of the spectro-temporal structure of sound. Internal models of these sound properties must play an essential role in optimizing proactive perceptual and cognitive performance. Speech, as a particularly complex sound signal, evolves over different timescales and requires spectral and temporal segmentation to establish building blocks for the construction of models of the auditory world. This fundamental task likely relies on the precise orchestration of cortical and subcortical brain areas (Kotz et al., 2014; Schwartze & Kotz, 2016). An integrative interpretation of the predicted results from the perspective of a cerebellar-temporal cortex interface - with potential lateralization reflecting differential temporal sensitivities - offers an intriguing new perspective to explore the functional basis of cerebellar internal modeling in motor control and audition, providing a computational generalization that may offer useful new angles for experimentation.

## Results

### Lesions and Behavioral Deficits

Twelve patients with chronic left temporal stroke and twelve matched controls (Tables S1-3) were mapped and tested using auditory temporal order and discrimination tasks (same-different judgments). We tested the groups on a range of perceptual tasks selected to probe temporal processing in hearing. Figures 1 and 2 depict the lesion distribution and task performance, respectively.

**Figure 1.**
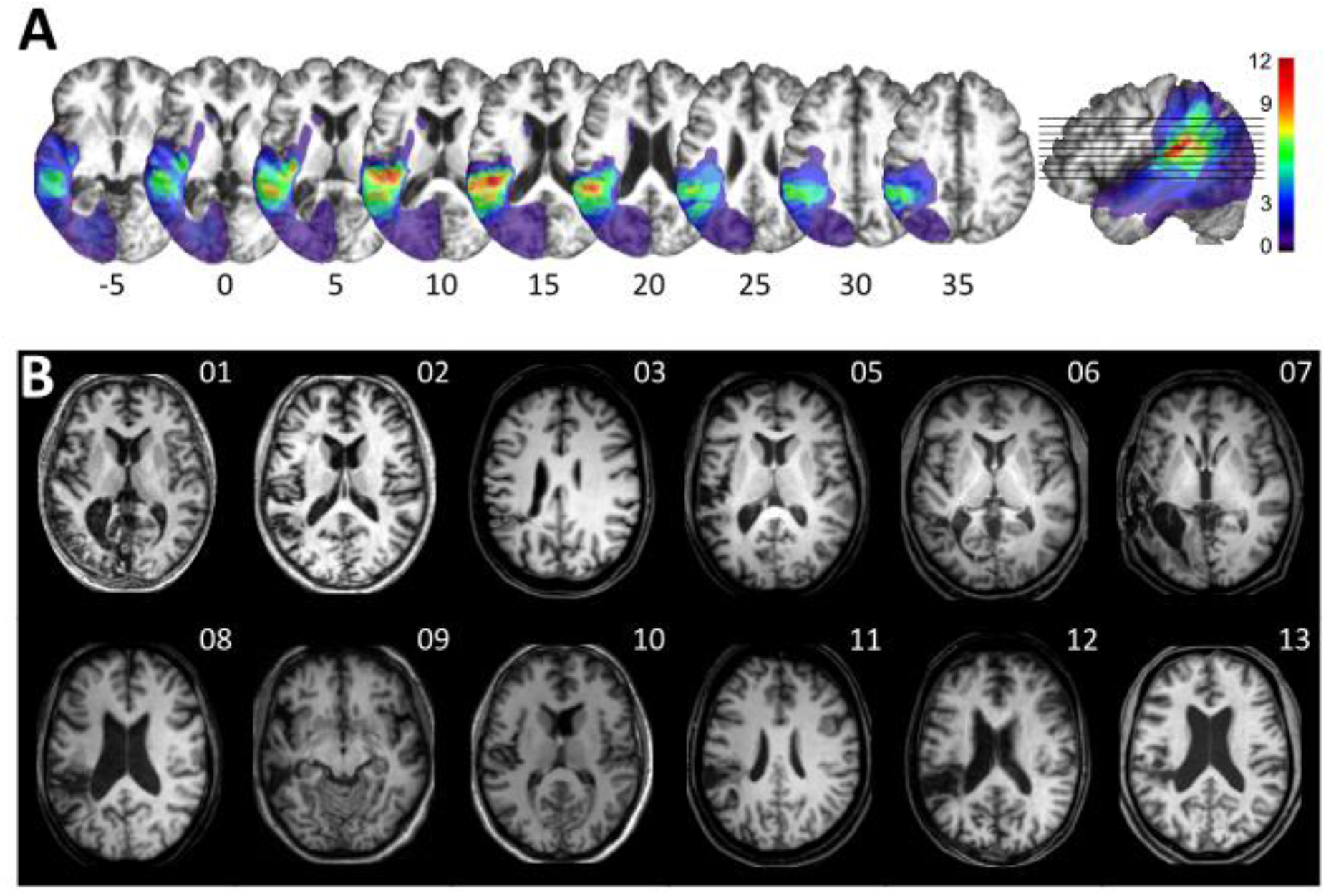
Visualization of lesion distribution. Data superimposed on the scalp-stripped mean patient T1-weighted image. Corresponding MNI coordinates for axial slices are shown on an orthogonal slice view (x = −45). (A) (top row) Lesion frequency map: lesion distribution in the twelve patients. Colorbar specifies number of patients with overlapping lesions in each voxel, with hot colors indicating that a greater number of patients had lesions in this region. Maximum lesion overlap in left posterior superior temporal gyrus (planum temporale) and underlying white matter (MNI −45, −36, 15).

**Figure 2.**
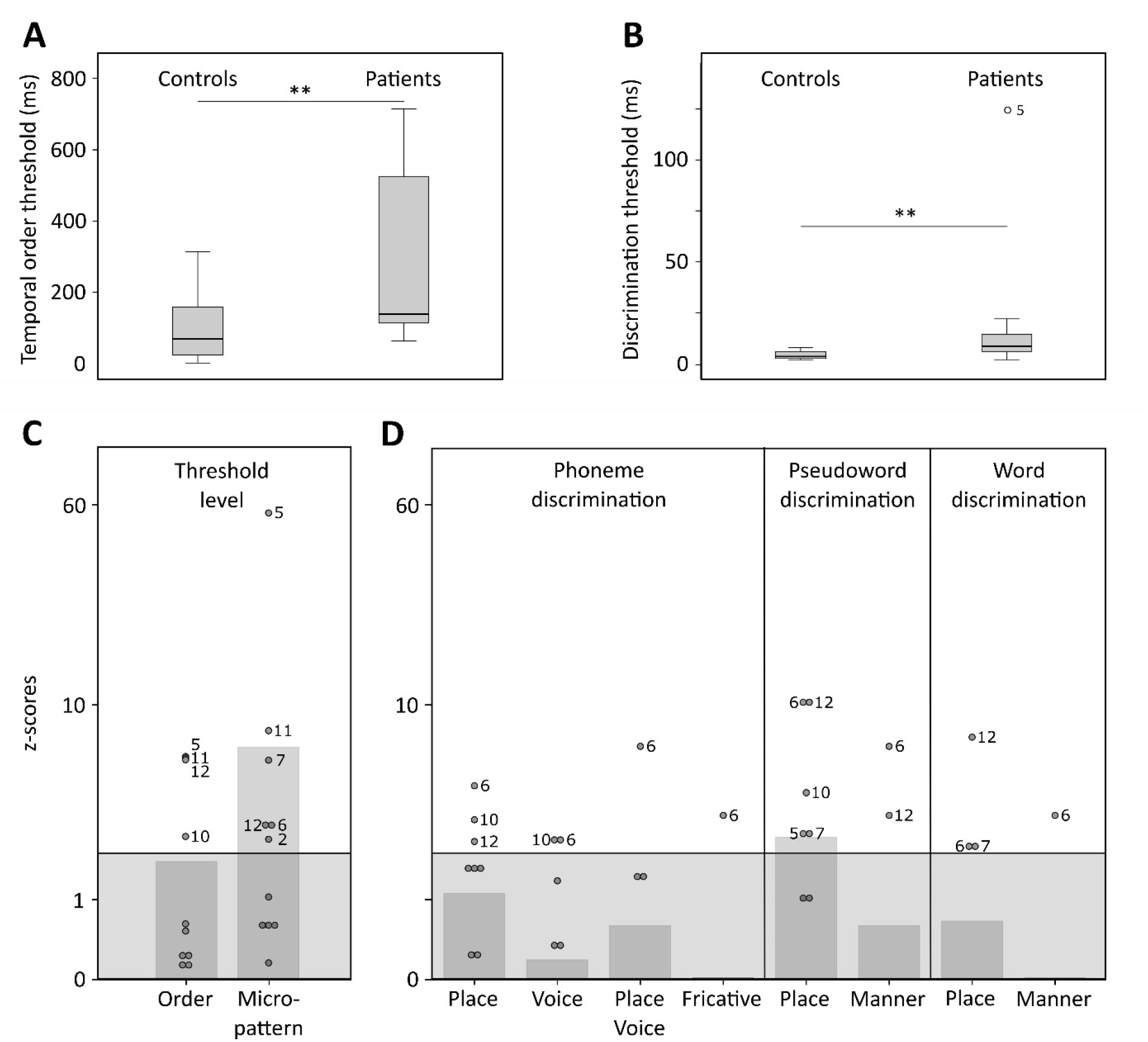
Temporal order and discrimination thresholds and identification of deficit positive (LG^+^) and negative (LG^-^) lesion group. Boxplots display median (horizontal line), first and third quartile (box), data range (wiskers) and outlier (dot) of the threshold levels in milliseconds for temporal order judgments (**A**) and discrimination of micropatterns (**B**) in the control and patient group. Patients as compared to controls show higher temporal order and micropattern discrimination thresholds. To identify deficit positive (LG^+^) and negative (LG^-^) lesion groups, patients mean (bars, dark grey) and individual performance (circles) on temporal order and micropattern discrimination (C) and phoneme/word discrimination (D) were converted to into z-scores relative to control group means for each behavioral test. Values > 0 indicate worse performance than controls within (light gray) and outside (no color) plus 2 standard deviations (SD) of the controls mean. Patients scoring outside 2 SD of the controls (impaired performance, LG^+^) are indicated by subject number.

Tonal stimuli were presented to determine individual threshold levels for temporal order judgment and micropattern discrimination, which were then linked to normal performance in controls (Figure 2). These tasks index a participant’s perceptual abilities to (i) encode non-verbal short time-scale spectral information, and (ii) compare a current stimulus to the representation of a preceding stimulus. The term ‘micropattern’ describes pairs of complex tones presented with stimulus onset asynchronies (SOA) below an individual’s temporal order threshold, i.e., the shortest SOA at which the temporal order of two tones can be perceived (Chedru et al., 1978; Pöppel, 1978). In this case, the order of still discriminable different stimulus elements cannot be determined. However, frequency reversals within a micropattern lead to a perceptual dissociation. Micropatterns are perceived as relatively lower or higher in pitch, a phenomenon attributed to the perceptual dominance of the second stimulus frequency (Efron, 1973).

Same-different judgments were used to assess the discrimination performance for minimal-pair words (example: Dach [dax] (engl. *roof*) – Bach [bax] (engl. *stream*)), non-words (example: Pach [pax] – Kach [kax]), and phonemes (example: e.g., /tr/ and /pr/) differing in contrastive phonological features. Corresponding error rates served to compare basic perceptual speech abilities for the encoding and modeling of different levels of speech and contrastive features (e.g., articulation, voicing, manner of articulation) relative to controls (Figure 2D). See Methods for details.

### Impaired Encoding of non-verbal and verbal spectro-temporal information

Healthy controls showed typical threshold values comparable to previously reported temporal order (SOAs above 15-60 ms) and micropattern discrimination tasks (SOAs of at least of 5 ms) (Efron, 1963, 1973; Hirsh & Sherrick, 1961; Yund & Efron, 1974). In contrast, patients required longer intervals to judge the order of two different frequency tones (U = 114, p = .007, effect size *r* = .495; Figure 2A and Table S4) and to discriminate two micropatterns (U = 120.5, p = .002, *r* = .572; Figure 2B and Table S4). Order and discrimination thresholds were positively correlated in patients (*p =* .015, Spearman’s Rho *r*_*s*_ = .681) and controls (*p =* .006, *r*_*s*_ = .735). Error rates differed in patients and controls for the discrimination of words, pseudowords, and phonemes (Friedman test, χ^2^(2) = 16.026, *p* = .0002). Phoneme discrimination displayed increased error rates compared to words (*Z* = - 3.624, *p* = .0003, *r* = .740, Bonferroni adjustment at *p* < .017) and pseudowords compared to words (*Z* = −2.684, *p* = .007, *r* = .548). Although healthy controls did not show any category effect, patients (χ^2^(2) = 10.857, *p* = .003) showed higher error rates for phonemes than words (*Z* = −2.287, *p* = .015, *r* = .660), and for pseudowords than words (*Z* = −2.666, *p* = .002, *r* = .769). Subsequent between-group comparisons revealed non-significant trends for higher error rates in patients compared to controls for the discrimination of pseudoword and phoneme pairs (Table 1). Thus, in line with previous results (Chedru et al., 1978; Efron, 1963; Swisher & Hirsh, 1972), we confirm that patients with left posterior temporal strokes show less accurate auditory spectro-temporal processing and concomitant perceptual speech deficits.

**Table 1.** Between-group comparisons of error rates for verbal discrimination tasks per category and contrastive feature. Mean relative error rates (±SD) and non-parametric test statistics for within (Friedman χ^2^ and Wilcoxon signed-rank Z-statistics, Bonferroni-adjusted significance levels set at p < .017) and between-subject comparisons (Mann-Whitney-U test, exact p-values (one-sided)) for each category and contrastive feature.

| Variable | Group |  | Test statistics |  |  |
| --- | --- | --- | --- | --- | --- |
| Category | Patients | Controls | Category effect |  | Group effects |
|  | (N = 12) | (N = 12) | Patients | Controls |  |
| Words (W) | .02 ± .02 | .01 ± .01 | $\chi^2(2) = 10.86$ | $\chi^2(2) = 5.72$ | U = 83, p = .276 |
| Pseudowords (PW) | .05 ± .05 | .02 ± .03 | p = .003 | p = .056 | U = 96, p = .089 |
| Phonemes (P) | .07 ± .09 | .04 ± .07 |  |  | U = 95, p = .099 |
| Feature | Patients | Controls | Feature effects |  | Group effects |
|  | (N = 12) | (N = 12) | Patients | Controls |  |
| Place (W) | .04 ± .08 | .01 ± .03 | Z = -1.34 | Z = -1 | U = 79, p = .366 |
| Manner (W) | .01 ± .02 | .01 ± .02 | p = .250 | p = .500 | U = 72, p = .500 |
| Place (PW) | .16 ± .22 | .03 ± .05 | Z = -2.38 | Z = -1.342 | U = 101.5, p = .045 |
| Manner (PW) | .02 ± .05 | .01 ± .02 | p = .008 | p = .250 | U = 78.5, p = .366 |
| Place (P) | .23 ± .23 | .08 ± .14 | $\chi^2(2) = 18.31$ | $\chi^2(2) = 7.0$ | U = 99.5, p = .057 |
| Fricatives (P) | .03 ± .12 | .03 ± .12 | p = .00004 | p = .065 | U = 72, p = .500 |
| Voice (P) | .07 ± .1 | .06 ± .08 |  |  | U = 70.5, p = .466 |
| Place and Voice (P) | .05 ± .15 | .03 ± .08 |  |  | U = 72.5, p = .500 |

### Feature Specific Impairment for Place of Articulation Contrasts

As the left-lateralized lesion patients showed robustly worse pseudoword and phoneme discrimination, we next tested phoneme specific features focusing specifically on the shortest time scales (Rosen, 1992). Only patients showed significant effects for contrastive features for phoneme (χ^2^(2) = 18.313, *p* = .00004) and pseudoword pairs (Z = −2.38, p = .008, *r* = .687). Post-hoc Wilcoxon tests (Bonferroni-adjusted significance level at *p* < .0083) confirmed higher error rates for the discrimination of place of articulation contrasts than for voicing (*Z* = - 2.521, *p* = .004, *r* = .728), combined place of articulation and voicing (*Z* = - 2.536, *p* = .004, *r* = .732), fricative (*Z* = - 2.555, *p* = .004, *r* = .738), and manner of articulation contrasts (*Z* = −2.684, *p* = .004, *r* = .775) in phonemes and pseudowords. The same tests only yielded a non-significant trend for contrastive feature within the phoneme category (χ^2^(2) = 7.0, *p* = .065) in healthy controls. Subsequent between-group comparisons revealed higher error rates for the discrimination of place of articulation contrasts in pseudowords in patients but not controls (*U* = 101.5, *p* = .045, *r* = .348) and a non-significant trend (*U* = 99.5, *p* = .057, *r* = .343) for the discrimination of place of articulation contrasts in phonemes (Table 1). In sum, patients displayed speech processing deficits preeminent for information encoded in the spectro-temporal fine structure at short time scales (Rosen, 1992).

### Linking Auditory Spectro-temporal and Short Time-scale Phonemic Processing

Threshold levels for temporal order and micropattern discrimination were correlated (Spearman’s rank-order correlations, one-sided) with performance measures for different discriminative features to assess possible associations between impaired auditory temporal processing and the processing of phonemic cues encoded at short time scales. There was a positive correlation between error rates for voicing contrasts in phoneme discrimination with threshold levels for auditory order (*p* = .032, *r* = .549) and micropattern discrimination (*p* = .043, *r* = .516). The associations between increased error rates for place of articulation contrasts with increased auditory order (*p* = .051, *r* = .495) and discrimination thresholds did not meet conventional significance (*p* = .061, *r* = .471).

Importantly, no significant associations were found between lesion volume or hearing loss and threshold values or error rates for feature discrimination (Table S5). In line with our hypotheses and previous findings (Fink et al., 2006; Robson et al., 2013), we show that *patients exhibit a short-time scale specific perceptual deficit* for tones and phonemes.

### Posterior Superior Temporal Injury is Associated with Impaired Spectro-Temporal Encoding

Despite consistent group-level effects, patients varied in terms of their encoding capacity for tones and phonemes. We therefore next explored whether performance variability links to lesion sites within the left temporal cortex. Subsequent analyses related deficient spectro-temporal processing at short timescales to specific temporal lesion sites (lesion-symptom mapping). Performance differences between healthy controls and patients with or without a performance impairment (see Figure 2) allowed identification of brain regions more frequently associated with an impairment (Figure 3; see Methods for more details). This procedure effectively split the patient group into two sub-groups of equal size, as six patients showed impaired performance in at least two subtests (deficit positive lesion group, LG^+^) as compared to six patients (deficit negative lesion group, LG^-^) performing similarly to controls (Figure 2C-D). The patient groups did not differ in terms of demographic or clinical characteristics (Table S6). The maximal lesion overlap (Figure 1A) was centered on the posterior superior surface of the left superior temporal gyrus (planum temporale) extending into the underlying white matter. In contrast, the lesion subtraction of LG^-^patients from LG^+^ patients (Figure 3A) linked posterior superior temporal sulcus (STS), adjacent middle temporal gyrus (MTG), and white matter below the STS to the spectro-temporal encoding impairment in the LG^+^ group. Differences in lesion distribution between LG^-^and LG^*^patients were statistically confirmed (Liebermeister test, binomial data with 1 = LG^+^ and 0 = LG^-^, permutation FWE-corrected α-level of *p* < 0.05) (Figure 3B).

**Figure 3.**
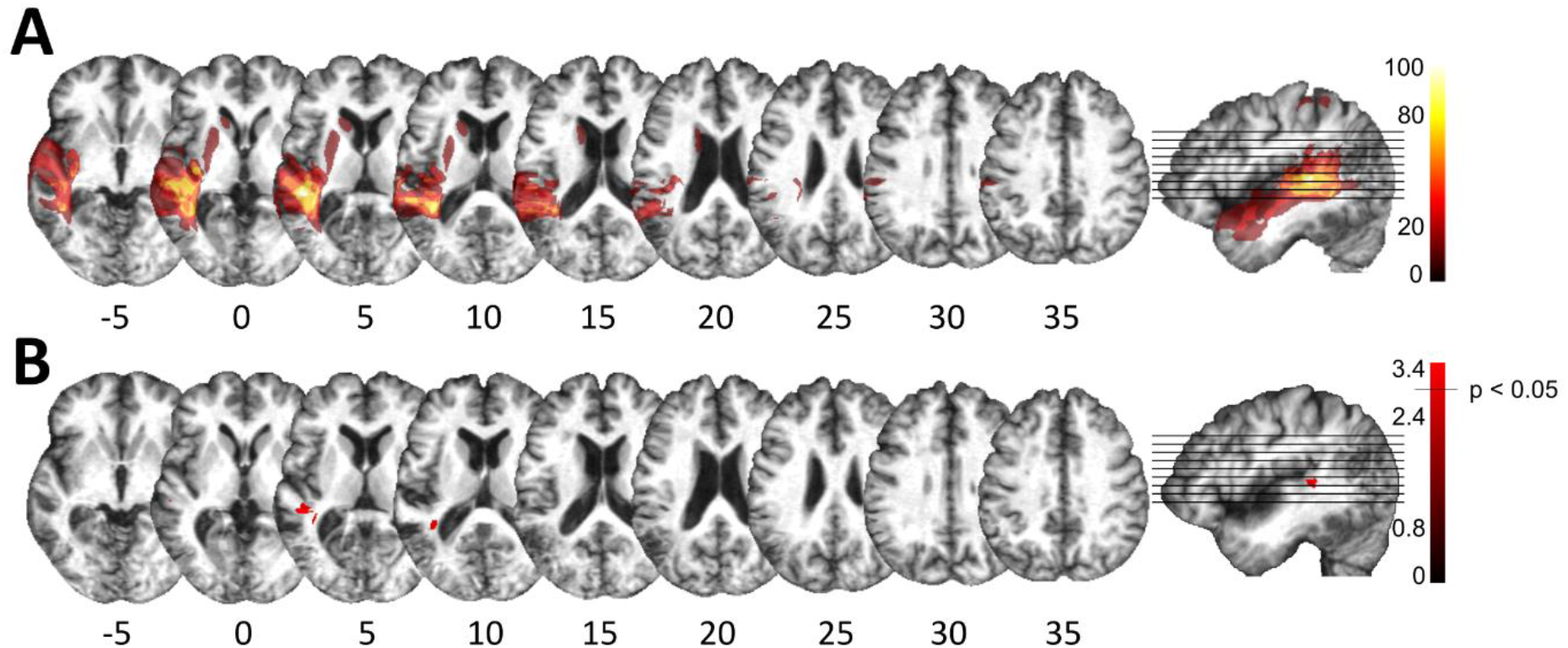
Lesion analysis of deficit positive (LG^+^) and negative (LG^-^) lesion group. (**A**) Subtraction plot shows voxels more frequently damaged in LG^+^. Colorbar specifies relative frequency (percentage) of overlapping lesions in the patient group with impaired performance (LG^+^) after subtracting lesion overlap of LG^-^from lesion overlap of LG^+^. (**B**) Voxelwise statistical analyses (Liebermeister measure for binomial data, permutation FWE-corrected z-scores at α-level of p < 0.05): lesions in posterior STS (MNI −48, −34, 5 and −38, −43, 10) are significantly associated with impaired temporal information processing (LG^+^).

### Posterior Superior Temporal Projections interface Spectro-Temporal Processing Networks with the Cerebellum

We next used the respective areas as seed regions for probabilistic fiber tractography in a healthy age-matched sample to visualize the underlying common connectivity pattern (see Methods). Long association and projection fibers originating from the seed masks in the posterior STS and the subjacent white matter below the STS (Figure 5A) were identified within the anterior floor of the left external capsule, the left periventricular white matter, and the brain stem. Subdivision into separate fiber bundles (Figure 4) indicated connectivity along the inferior fronto-occipital fasciculus (IFOF) travelling within the anterior floor of the external capsule. Terminations were present in the left inferior frontal gyrus (pars triangularis, Brodmann area (BA) 45) and in the lateral orbitofrontal cortex (BA 47). Association fibers also extended along the posterior lateral surface of the lateral ventricle to the left superior parietal cortex (BA 7). Considering the characteristic connectivity patterns of these association fibers with the superior parietal cortex, they may correspond to the posterior middle longitudinal fasciculus (Makris, Preti, Asami, et al., 2013; Wang et al., 2013). Other cortico-cortical association fibers curved upward and travelled rostrally within the periventricular white matter lateral to the corona radiata in the superior longitudinal fasciculus (SLF) with terminations in the left inferior frontal gyrus (pars opercularis, BA 44), and the left dorsolateral prefrontal cortex (BA 9, BA 46).

**Figure 4.**
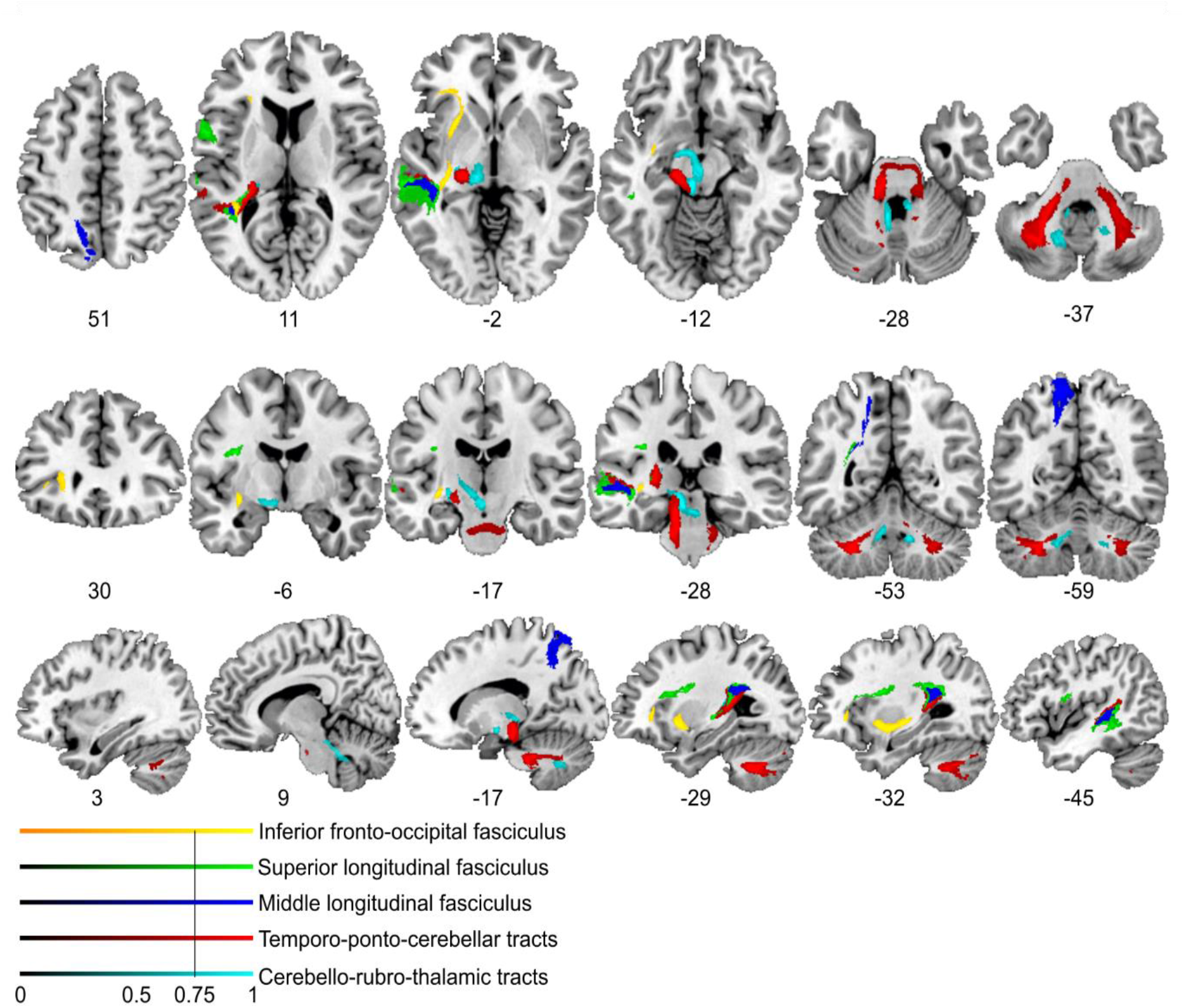
Lesion-informed probabilistic tractography. Diffusion tractography based on a dataset of 12 healthy controls. Seed areas only included voxels being more frequently associated with impaired processing of temporal information. Inclusion masks were used to subdivide individual connectivity distributions into separate fiber bundles. The tracts are superimposed on the MRIcron ch2bet template in standard MNI space (axial, coronal and sagittal slices, corresponding MNI coordinates are indicated below). Displayed group variability maps result from binarized tract volumes (thresholded connectivity distributions) that quantify the percentage of subjects (> 75 %) showing connectivity between the seed masks and the respective voxel (values range from 0.0 – 1.0). Yellow: Inferior fronto-occipital fasciculus (IFOF), green: superior longitudinal fasciculus (SLF), red: temporo-ponto-cerebellar tracts, dark blue: middle longitudinal fasciculus, light blue: cerebello-rubro-thalamic tract.

**Figure 5.**
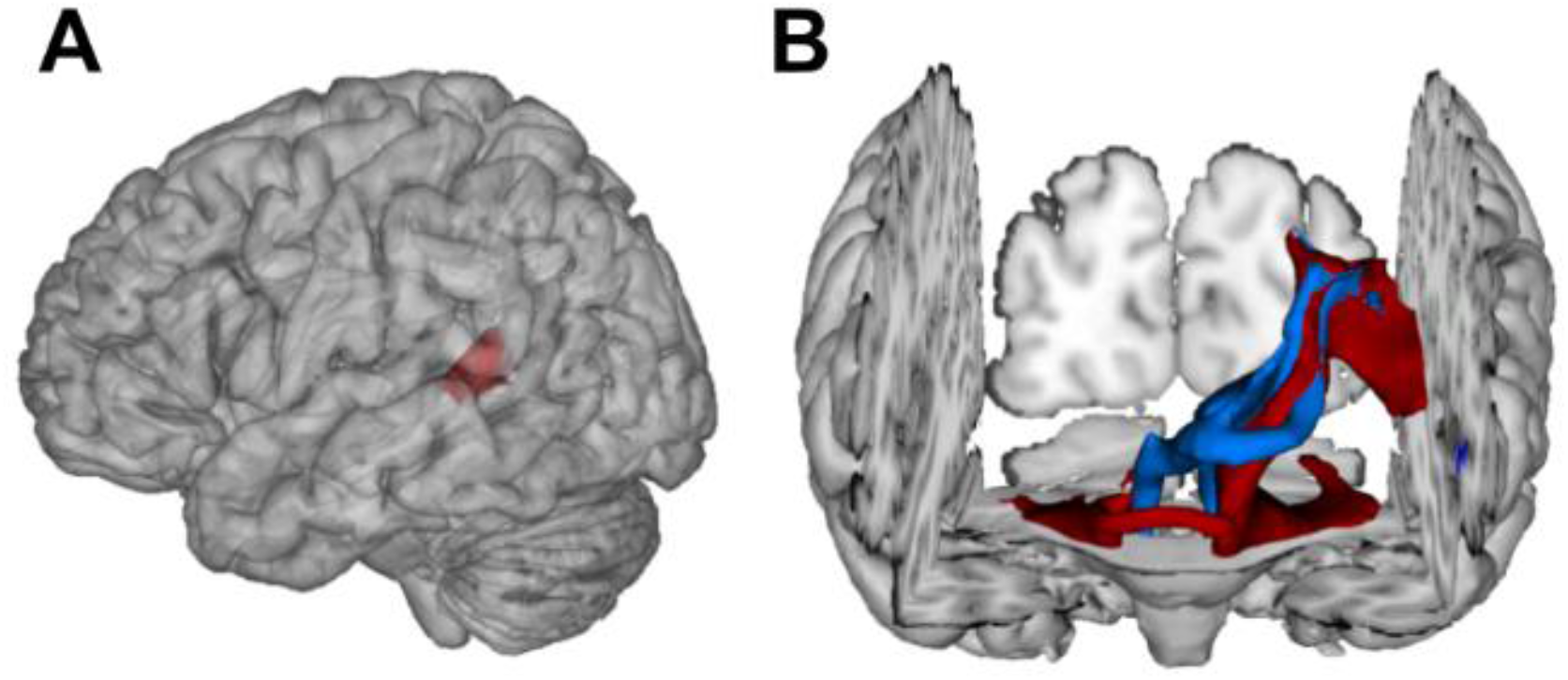
Visualization of temporal cortex-cerebellum connectivity. Bilateral and bidirectional connectivity of seed regions (**A**) in the left posterior superior temporal sulcus (pSTS). (**B**) Temporo-ponto-cerebellar tracts (red) and cerebello-rubro-thalamo-temporal tracts (blue) connect pSTS with the postero-lateral cerebellum and the dentate nucleus, respectively.

Most importantly, fibers originating in the superior posterior temporal cortex reached the posterolateral cerebellum (crura I and II) and the dentate nuclei bilaterally (Figures 4 and 5B). These fibers coursed rostrally and ascended medially in close proximity to the posterior temporal and inferior parietal cortex before they descended through the retrolenticular internal capsule along the left cerebral peduncle to the pontine nuclei and the ipsilateral middle cerebellar peduncle. They additionally decussated at the ventral pons to the right cerebellar peduncle, giving rise to bilateral temporo-ponto-cerebellar tracts (Figure 5B). Other fibers conformed to the cerebello-rubro-thalamic tract connecting the posterior superior temporal cortex with the bilateral dentate nuclei along the superior cerebellar peduncles. They crossed at the level of the inferior olive to the left red nucleus and projected to the posterior thalamus (pulvinar) (Figures 4 and 5B). Although the cerebello-rubro-thalamic tract is considered a decussating pathway, there is some evidence for a non-decussating pathway and for specific connections of these pathways to more anterior and lateral as opposed to more posterior and medial thalamic targets (Petersen et al., 2018). Considering that the differential connectivity of Broca’s area with the thalamus includes the pulvinar, one may speculate that the non-decussating pathway also supports language function (Bohsali et al., 2015).

Taken together, the results demonstrate that lesions in the left posterior STS led to a spectro-temporal processing deficit at short timescales for lower level auditory and speech information. The neural substrate for such perceptual necessities, is revealed here for the first time: a bidirectional temporo-cerebellar connectivity revealed by probabilistic tractography.

## Discussion

Our functional anatomic discovery, deriving from deficit lesion data from patients as well as tractography data from healthy participants, provides a mechanistic link between two domains of inquiry that have proceeded largely independently: research on the temporal dynamics of auditory perception and research on internal models supported by the cerebellum. The newly described cortex-cerebellum connectivity forms the basis for how the specific anatomic layout underpins one key aspect of auditory perceptual analysis. We set out to answer the following questions: First, how does the asymmetrical sampling of verbal and non-verbal sound information at different timescales tie in with the encoding of spectro-temporal structure in an internal model framework? Second, do we have to consider cross-lateral cortico-subcortical structural connectivity to achieve a comprehensive view of asymmetrical sampling of sound properties? Lesion-symptom informed probabilistic tractography, seeded in the left posterior STS of healthy participants, revealed temporo-frontal and bidirectional structural connectivity with the cerebellar dentate nuclei and crura I/II (see also Sokolov, Erb, Grodd, and Pavlova (2014) for right pSTS connectivity). The evidence we describe (i) supports the hypothesis that encoding and modeling of spectro-temporal information relies on a temporo-cerebellar system and (ii) suggests a generalizable role of cerebellar-mediated internal models that extends beyond motor control to auditory perception.

### Impaired auditory spectro-temporal encoding for non-verbal and verbal information

The tested patient sample (see Figure 1) displayed only mild aphasic symptoms (Table S3) but higher temporal order thresholds (Figure 2A), falling into the range of 150-600 ms relative to 15-60 ms previously reported for healthy participants (Efron, 1963; Fink et al., 2006). Higher micropattern discrimination thresholds in patients (Figure 2B) converge with evidence for impaired micropattern discrimination in patients with selective temporal compared to frontal lesions (Chedru et al., 1978). In patients, discrimination of the place of articulation contrast was impaired for phonemes and pseudowords but not words (Figure 2D). This feature is likely represented in the spectro-temporal fine structure occurring over short timescales (20-50 ms), whereas phonemic contrasts to voicing and manner are encoded in time-scale duration differences of voice-onset times (Elangovan & Stuart, 2008) and the slowly varying temporal envelope (50-500 ms; Rosen, 1992). The redundancy of information across shorter and longer timescales for the latter phenomena may contribute to this feature specific effect. Both phoneme discrimination and auditory order or micropattern discrimination are assumed to map onto lower-level auditory processes on short timescales. In contrast to earlier findings (Chedru et al., 1978), we show that discrimination thresholds and discrimination performance for non-verbal and verbal information go hand in hand. This suggests a common process that contributes to short timescale spectro-temporal encoding for both non-verbal *and* verbal information - and that impaired auditory temporal processing can (at least partly) explain lower-level auditory deficits cascading into speech comprehension deficits (Robson et al., 2013).

### Posterior Superior Temporal Regions Link Cortico-cortical and Subcortico-cortical Spectro-temporal Processing Networks

Lesion-symptom mapping (Figure 3) identified an area in the posterior superior temporal sulcus (pSTS) that was more frequently affected in patients with impaired short timescale spectro-temporal encoding for non-verbal and verbal information. This is in line with the dissociation of information unfolding over timescales corresponding to global prosodic, syllabic, and phonemic levels that is mirrored by a functional asymmetric temporal sensitivity of higher-order auditory areas in the superior temporal cortices. These are thought to preferentially encode rapidly changing auditory signals in time windows of ∼20-40 ms in the left hemisphere and of ∼150-250 ms in the right hemisphere (*asymmetric sampling in time;* Boemio et al., 2005; Flinker et al. 2019; Poeppel, 2003).

Our tractography findings of *cortico-cortical structural connectivity* are in line with results mapping speech processing networks that connect to the left MTG/STS along the inferior-fronto-occipital-fasciculus (IFOF) and the superior longitudinal fasciculus (SLF) with BA 47 and BA 46 (Turken & Dronkers, 2011). Functionally distinct subdivisions of the SLF, namely the SLF III and arcuate fasciculus, may provide higher order (somato-)sensory and auditory input to inferior frontal (BA 44) or dorsolateral prefrontal regions (BA 46, BA 6/8; Makris et al., 2005). Other studies revealed connectivity along the middle longitudinal fascicle between the posterior temporal cortex and the superior parietal lobe (BA 7; Makris et al., 2013; Wang et al., 2013), an area associated with audio-visual multisensory integration (Molholm et al., 2006).

We add a novel contribution to this established pattern of cortico-cortical connectivity by revealing cross-lateral structural cortico-subcortical structural connectivity, with clear implications for the emerging cerebellar contributions to higher cognitive functions. To date, such contributions have been related to reciprocal prefrontal-cerebellar and posterior parieto-cerebellar projections, connecting association cortices with the lateral and posterior cerebellar hemispheres (Brodal, 1978a, 1978b, 1979; Jissendi, Baudry, & Baleriaux, 2008; Kelly & Strick, 2003; Ramnani et al., 2006). Earlier claims considering temporo-cerebellar projections based on diffusion-weighted imaging of the cerebral peduncles as insignificant in both humans and non-human primates (Ramnani et al., 2006), should be revised, as the topographical distribution of cortico-ponto-cerebellar projections likely extends beyond the most prominent prefrontal and primary motor structural connectivity patterns. As demonstrated here, the cortico-cerebellar system may also comprise reciprocal ipsi- and contralateral temporo-cerebellar projections that so far have gained only very little attention (Schmahmann & Pandya, 2009; Schmahmann & Pandya, 1991; Sokolov et al., 2014). Moreover, the present findings confirm fibers connecting the superior posterior temporal cortex and the posterior lateral cerebellum (crura I/II, cerebello-rubro-thalamic tract) and indicate fibers originating in bilateral dentate nuclei, which run along the superior cerebellar peduncle and posterior thalamus (cerebello-rubro-thalamic tract). These observations confirm the concept of reciprocal cortico-cerebello-cortical loops (Salmi et al., 2010) and conform to anatomical landmarks for cerebro-cerebellar connections in the brainstem and thalamus demonstrated in previous neuroanatomical and MRI studies (Bernard et al., 2014; Brodal, 1978a, 1979; Dum & Strick, 2003; Habas & Cabanis, 2006, 2007; Pandya, Rosene, & Doolittle, 1994; Schmahmann & Pandya, 1991).

Consequently, the need arises to integrate these novel structural connectivity findings (i) with well-established functional evidence for spectro-temporal sound processing at different time scales in temporal cortices (Boemio et al., 2005; Callan et al., 2007; Poeppel, 2003), and (ii) with the cerebellum’s critical role in supporting internal models (Kotz & Schwartze, 2010; Schwartze, Tavano, Schröger, & Kotz, 2012).

### The temporo-cerebellar interface is linked to spectro-temporal encoding

In contrast to the bilateral posterior lateral cerebellar contributions to auditory (Pastor et al., 2002; Petacchi et al., 2005) and temporal processing (Ivry et al., 2002; Keele & Ivry, 1990; Spencer & Ivry, 2013), there is sparse evidence on temporo-cerebellar coupling (Pastor et al., 2002; Pastor, Thut, & Pascual-Leone, 2006; Pastor, Vidaurre, Fernandez-Seara, Villanueva, & Friston, 2008). However, such coupling may support the continuous updating of internal models of the auditory environment (Kotz et al., 2014) based on precise encoding of temporal structure. For example, functional imaging studies show increased induced oscillatory activity in response to 40 Hz auditory stimulation (corresponding to a sampling period of ∼25 ms) in the bilateral posterolateral cerebellum (crus II) (Pastor et al., 2002), and effective connectivity between superior temporal areas (STG and STS) and cerebellar crus II increases with 40 Hz auditory stimulation (Pastor et al., 2008). Likewise, cortical oscillatory responses to 40 Hz auditory stimulation diminish after inhibitory cerebellar transcranial magnetic stimulation (Pastor, Thut, et al., 2006). Moreover, induced gamma band oscillatory activity in the auditory cortex as well as in the cerebellum in response to random auditory stimulation tightly time-locks to auditory stimulus onsets (Demiralp, Basar-Eroglu, & Basar, 1996).

The cerebellum likely supports the encoding of an event-based representation of the temporal structure extracted from the auditory input signal by tracking salient modulations (e.g., changing sound energy levels as in onset transients or frequency transitions) of the physical sound across time that contribute to the segmentation of a continuous auditory input signal into smaller perceptual units. Such reciprocal temporo-cerebellar interactions might provide a unitary stimulus representation at different time scales for later processing stages (Schwartze & Kotz, 2016; Schwartze et al., 2012; Weise, Bendixen, Muller, & Schröger, 2012). The overlap with the encoding of auditory signals in temporal integration windows of different lengths (Boemio et al., 2005) and possible cerebellar encoding of event boundaries across different time-scales (Callan et al., 2007) suggests that a temporally structured event representation may map the detailed cortical representation of auditory sensory information to relevant points in time (Fig. 6) allowing for salient sound features to be optimally processed (Kotz & Schwartze, 2010; Schwartze & Kotz, 2013, 2016; Schwartze et al., 2012).

**Figure 6.**
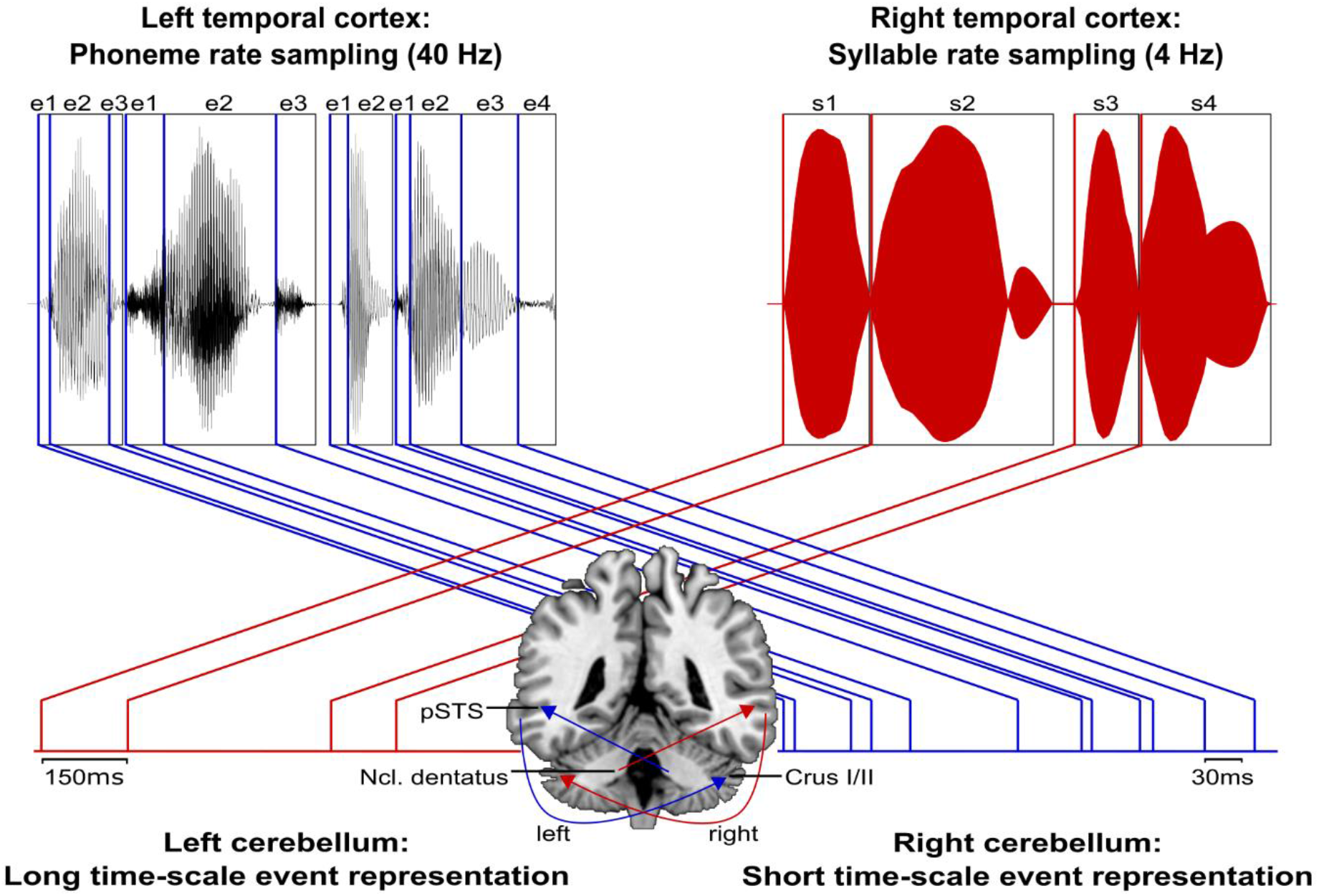
Schematic visualization of temporo-cerebellar interaction for internal model construction in audition. Left and right cerebellum contribute to the encoding of event boundaries across long (red) and short (blue) timescales, respectively (Callan et al., 2007). These event representations are extracted from salient modulations of sound properties, i.e., changes in the speech envelope (fluctuations in overall amplitude, red) corresponding to syllables (s1-s4) and the fine structure (formant frequency transitions, blue) corresponding to phonemes (e1-e4) (Rosen, 1992; Weise et al., 2012). Reciprocal temporo-cerebellar interactions between posterior superior temporal sulci (pSTS), crura I/II and dentate nuclei yield unitary temporally structured stimulus representations conveyed by temporo-ponto-cerebellar and cerebello-rubro-thalamo-temporal projections (arrows). The resulting internal representation of the temporal structure of sound sequences, e.g., speech, fits the detailed cortical representation of the auditory input to relevant points in time to guide the segmentation of a continuous input signal (waveform) into smaller perceptual units (boxes). This process allows distinctive sound features (e.g., word initial plosives /d/ (e1 in s1), /t/ (e1 in s2) and /b/ (e1 in s3) varying in voicing or place of articulation) to be optimally integrated at the time of their occurrence.

Notably, in auditory processing the prefrontal cortex (supplementary motor area, SMA, BA 6; (Pastor, Macaluso, Day, & Frackowiak, 2006)) and its connections to the cerebellum (Aso, Hanakawa, Aso, & Fukuyama, 2010; Jissendi et al., 2008) play an essential role in the temporal integration of sensory information (Schwartze & Kotz, 2013; Schwartze et al., 2012). This extended network consisting of prefrontal cortex, temporal cortex, and cerebellum provides a platform to integrate sensory input over different time scales in order to continuously update spectro-temporal models of the auditory environment, thus optimizing sound processing (Kotz & Schwartze, 2010; Kotz et al., 2014; Schwartze & Kotz, 2013; Schwartze et al., 2012).

Many investigations have put forward theories on the cerebellum’s role in perception and cognition (for a review, see (Baumann et al., 2015)), and the illustrated cortico-subcortical and cortico-cortical structural connectivity pattern is presumably not unique for auditory temporal processing but showcases one possible general role for perceptual processing.

## Conclusions

We show that left posterior temporal cortex is contributing to audition in a time-sensitive manner. This functional characteristic, identified by lesion symptom mapping, has led to the discovery of a specific cortico-subcortical structural connectivity pattern. Taken together, these results provide compelling evidence for a mechanism that in its simplicity not only applies to audition but may extend to other modalities relying on similar structural connectivity patterns (visual motion perception (Sokolov et al., 2014); multisensory integration (Baumann & Greenlee, 2007; Schmahmann & Pandya, 1991)). This points to a possible common neurobiological function (Schmahmann, 2004) supporting internal modeling of a dynamic environment.

## Materials and Methods

### Participants

A group of stroke patients (lesion group, LG, *N* = 12) and a group of healthy controls (control group, CG, *N* = 12) closely matched for handedness, gender, age and formal education (Table S1) were selected from databases at the Leipzig University Hospital Day Clinic for Cognitive Neurology and the Max Planck Institute for Human Cognitive and Brain Sciences, Leipzig, Germany. For patients (Table S2), initial inclusion criteria were a chronic ischemic stroke (time since lesion ≥ 12 months) in the left temporal lobe, no previous cerebral infarctions or lesions to other areas of the brain, and no history of other neurological or psychiatric disorders. To confirm normal hearing for their respective ages, patients and controls underwent audiometric screening with air conduction inside a sound-proof cabin (according to the *Guidelines for Manual Pure-Tone Threshold Audiometry*, American Speech-Language-Hearing Association, http://www.asha.org/docs/html/GL2005-00014.html) using a computer-based audiometer (MAICO MA 33, MAICO Diagnostic GmbH; headphones MAICO DD45; audiometric test frequencies 125 Hz to 8 kHz). One patient had to be excluded, as hearing thresholds between 0.5 and 2 kHz did not meet the criteria for age-normal hearing (International Standard ISO 7029, 2000). Hearing loss (dB HL) on both ears did not differ significantly between the remaining patients and the healthy controls (*0*.*5 kHz*: patients mean ± SD = 10.6 ± 3.7, controls = 12.3 ± 4.5 dB HL, U = 56, *p* = .38; *1*.*0 kHz*: patients = 9.4 ± 3.0, controls = 11.7 ± 5.6 dB HL, U = 55.5, *p* = .35; *1*.*5 kHz*: patients = 11.9 ± 4.0, controls = 13.3 ± 8.3 dB HL, U = 70.5, *p* = .93; *2 kHz*: patients = 12.9 ± 7.7, controls = 11.5 ± 7.0 dB HL, U = 78.5, *p* = .71). All participants were German native speakers and right-handed (handedness index score > 40). Time since lesion varied from 12 to 150 months (*M* = 60, *SD* = 42.1 months). Stroke severity was assessed by means of the National Institute of Health Stroke Scale (NIHSS, http://www.nihstrokescale.org/; German translation and validation (Berger et al., 1999)). The NIHSS (score range 0 – 42) provides a simple protocol to assess stroke-related motor (e.g., paralysis), non-motor and cognitive functions (e.g., consciousness, presence of aphasia or neglect). Although typically applied to patients suffering from acute stroke, the overall NIHSS scores for the current (chronic) patient group (*M* = 1.8 ± 1.0, range 0 – 4) reflect the severity of residual neurological deficits, ranging from normal function (score = 0) to minor impairments (scores = 1 – 4). However, NIHSS test items for language functions may not capture some residual deficits in chronic stroke patients. Therefore, language functions were assessed using the Aachener Aphasie Test (AAT) (Huber, Poeck, & Willmes, 1984) (Table S3).

For structural connectivity analysis, an additional group of 12 healthy participants was selected from the same databases. Individual patient-control pairs were matched in terms of age (51.8 ± 9.25 years), gender (7 male) and handedness (handedness index 83.8 ± 15.5; Edinburgh Handedness Inventory; (Oldfield, 1971)), to control for potential effects of these factors on fractional anisotropy (FA) values (Pal et al., 2011; Powell et al., 2012; Salat et al., 2005).

All experimental procedures were approved by the local ethics committee of the University of Leipzig according to the Declaration of Helsinki and written informed consent was given by each participant. All participants were naïve to the objective of the experiments and were financially compensated for their time and travel costs.

### Stimuli and Tasks

Speech stimuli (word, non-word, and phoneme pairs) were spoken in a soundproof cabin by a professionally trained female German native speaker, digitized with 16-bit resolution at a sampling rate of 44.1 kHz stereo using AlgoRec™ 2.1 (Algorithmix GmbH, Waldshut-Tiengen, Germany) and subsequently converted to mono. Offline editing comprised cutting at zero crossings before and after each word, normalization to an average intensity of 70 dB using the PRAAT software (www.praat.org; (Boersma & Weenink, 2001)). Non-speech stimuli were synthesized using Audacity® (www.audacity.sourceforge.net). The complex tones consisted of two 1000 and 2000 Hz (Δf = 1000 Hz) sinusoidal components with an average intensity of 70 dB and rise-plateau-decay values of 7-ms-10-ms-7-ms (Wright, 1960). These parameters and the stimulus presentation mode are based on previous work and allow for threshold determination in clinical populations (Fink, Churan, & Wittmann, 2005). All speech and non-speech stimuli were presented binaurally at a fixed intensity level via headphones (Sennheiser HD 202).

Individual threshold levels for *auditory temporal order judgments* (task 1) and *discrimination of complex tones (micropatterns)* (task 2) were determined to evaluate participants’ perceptual abilities to process non-verbal auditory spectro-temporal information. For task 1, auditory temporal order thresholds were defined as the shortest stimulus-onset-asynchrony (SOA) at which participants are able to correctly judge the temporal order of the two consecutively presented tone components (component A following B or B following A). For task 2, auditory discrimination thresholds referred to the shortest SOA at which participants are able to judge the difference between two complex tones (AB sounds perceptually different from BA but identical to AB). Participants were familiarized with the tones during a training period to ensure that they were readily audible to allow for order and perceptual difference judgments to be obtained. Participants completed a forced-choice decision for both tasks with the brief randomly presented complex tones, separated by SOAs ranging from 1000 ms (suprathreshold level for all participants) to 2 ms. For auditory order threshold determination (task 1) participants indicated whether the pitch of the first was higher or lower than the second component of the complex tone. A 50 ms-down 5 ms-up fixed step size staircase procedure with an SOA decrease after three correct and an increase after an incorrect response (reversal) was used. For discrimination thresholds (task 2) the complex tones were randomly presented below the individual temporal order threshold as determined by the previous test. Each presentation consisted of two complex tones in which the components had either the same (e.g., AB - AB) or reversed temporal order (e.g., AB - BA). Participants indicated whether the two complex tones were perceived as same or different. The same staircase procedure was applied, using fixed step sizes of 20 ms-down and 2 ms-up. The tasks were terminated after eight staircase reversals and auditory order (task 1) and discrimination (task 2) thresholds were calculated as the average of the last five SOAs at the staircase reversals points. This led to a probability of a 79.4 % correct threshold level for auditory discrimination and order judgments (Levitt, 1971).

To evaluate basic receptive and discriminative language abilities in the two groups, we used a test for auditory discrimination of words and non-words taken from the German LeMo Test battery (German version of *<u>Le</u>xikon <u>mo</u>dellorientiert*, model-based assessment of aphasia; (De Bleser, Cholewa, Tabatabaie, & Stadie, 1997; De Bleser, Stadie, Cholewa, & Tabatabaie, 2004). Participants had to judge whether spoken mono-morphemic minimal pairs (in two separate word and pseudoword lists) were identical or not (e.g., words: Dach [dax] (engl. *roof*) – Bach [bax] (engl. *stream*); pseudowords: Pach [pax] – Kach [kax]). Each list consisted of 72 items, 36 of which were identical. Non-identical items varied in terms of their consonantal features in either place (e.g., Bauk [bauk] – Baup [pauk]) or manner (e.g., Korf [korf] – Korm [korm]) of articulation. Pseudoword and word lists were presented separately to each participant.

Further language testing involved consonant feature related *discrimination of phoneme pairs* (unpublished material, Day Clinic for Cognitive Neurology, Leipzig). A total of 56 consonant clusters (e.g., /tr/ and /pr/) or single consonants (e.g., /t/ and /p/) that were either identical or varied in terms of their contrastive features were presented. Upon auditory presentation participants were asked to judge whether the presented pairs were the same (21 items) or different (35 items). Phonemic contrasts of non-identical pairs resulted from differences either in place of articulation (e.g., /p/ and /t/), voicing (e.g., /p/ and /b/), place of articulation and voicing (e.g., /p/ and /g/) or were fricative contrasts (e.g., /f/ and /ʃ/).

A short familiarization period consisting of five additional word, pseudoword, or phoneme pairs preceded the testing phase. Feedback was only provided during the training period and items were not repeated during the testing phase.

### Data acquisition

#### Structural and Diffusion weighted imaging

High-resolution anatomical T1 and T2-weighted magnetic resonance (MR) scans suitable for lesion reconstruction were available for all patients. Structural and diffusion-weighted data sets used for probabilistic tractography in healthy elderly participants were acquired with standard imaging protocols (see Imaging Procedures in Supplemental Information for scan parameters and processing pipeline).

### Data Analysis

#### Language and Behavioral Data

All statistical analyses were performed using SPSS 20.0 (IBM Corp., 2011). Means and standard deviations were calculated for the respective variables and groups. Shapiro-Wilk normality tests revealed that most of the data violated the assumption of normality for any of the psychometric variables. Therefore, non-parametric test methods were applied. Exact p-values (α = .05) are reported for small sample sizes. For independent samples one-sided Mann-Whitney-U tests were used to examine whether error rates or threshold levels were significantly higher in patients than in controls. Friedman tests with subsequent post-hoc Wilcoxon-signed rank tests (two-sided) were performed to test for within-group differences in error rates between linguistic stimulus categories (words, pseudowords and phonemes) or contrastive features (place or manner of articulation for words and pseudowords; place, voicing, place and voicing or fricatives for phonemes). Bonferroni-adjustment was applied if necessary. Associations between different linguistic and non-linguistic stimulus categories were analyzed by means of one-sided Spearman’s rank-order correlations coefficient in order to test for an increasing (positive) relationship between error rates and threshold levels in patients. Categorical variables were analyzed with Fisher’s exact test.

#### Lesion Mapping and Subtraction

Individual lesions were manually delineated on axial slices (slice thickness 1 mm) of T1-weighted images. To improve lesion characterization, lesion mapping was guided by co-registered T2-FLAIR images. The MRIcron software (http://www.mccauslandcenter.sc.edu/mricro/mricron/) was used to create a binary lesion map (volume of interest, VOI) for each subject, to generate individual lesion masks for normalization, and to estimate lesion volumes (Table S2). Individual T1-weighted images, co-registered T2-FLAIR and lesion maps were spatially normalized to standard stereotaxic Montreal Neurological Institute (MNI) space by means of the unified segmentation approach (Ashburner & Friston, 2005; Crinion et al., 2007) using *Clinical toolbox* (Rorden, Bonilha, Fridriksson, Bender, & Karnath, 2012) in SPM8 (Wellcome Department of Imaging Neuroscience, London, http://www.fil.ion.ucl.ac.uk/spm). This toolbox provides an MRI template and spatial priors for elderly participants. Cost-function masking was applied during normalization to achieve optimal anatomical co-localization of brain structures (Andersen, Rapcsak, & Beeson, 2010). All neuroanatomical data related to individual lesions, as well as lesion overlap and subtraction plots are reported in MNI space. Anatomical specification was based on visual inspection along with the macro-anatomical labels provided by the SPM based Anatomy Toolbox (Eickhoff et al., 2005). For visualization of overall lesion distribution individual lesion maps were superimposed (lesion overlap) on the scalp-stripped mean patient T1-weighted image (SPM8, ImCalc). Subsequent subtraction analysis was aimed at linking language and behavioral deficits related to representation of auditory temporal information to the anatomy of brain tissue damage (lesion-symptom mapping). These subtraction analyses account for differences between brain regions specifically contributing to a certain function and more vulnerable, but commonly damaged ones. This is achieved by contrasting lesions of patients with and without a (behavioral) deficit of interest at a certain behavioral cut-off value (Liebermann, Ploner, Kraft, Kopp, & Ostendorf, 2013; Rorden & Karnath, 2004; Rorden, Karnath, & Bonilha, 2007). Raw data from each subtest were transformed into z-scores corresponding to the controls mean and SD to group patients by deficit (impaired performance) based on behavioral data (Liebermann et al., 2013). Patients performing outside two SD of the healthy control groups mean in at least two of the speech (discrimination of word, pseudoword and phoneme pairs) or non-speech (auditory order and MP discrimination thresholds) subtests were assigned to the group with impaired performance (deficit positive lesion group, LG^+^). Remaining patients were assigned to the control patient group performing within the normal range (two SD of the control groups mean) on speech and non-speech subtests in the same set of experiments (deficit negative lesion group, LG^-^). Groups (LG^+^ and LG^-^) were compared for differences in age, gender, handedness, education level, lesion volume, time since lesion and performance on the Token test using independent sample t-tests. Subtraction plots were estimated from individual VOIs using MRIcron (http://www.mccauslandcenter.sc.edu/mricro/mricron/). The resulting subtraction plots indicate the relative frequency (percentage) of overlapping lesions in the patient group with abnormal performance (LG^+^) after subtracting the lesion overlap of LG^-^from the overlap of LG^+^. To infer statistical significance of differences in lesion distribution between LG^+^ and LG^-^, the non-parametric Liebermeister test was applied to the binominal data that reflected group classification. Non-parametric mapping (NPM, distributed with MRIcron) was used for voxelwise statistical analysis including only voxels affected in at least one patient and correcting for multiple comparisons by permutation testing (N = 12, permutations = 4000) (Rorden et al., 2007).

#### Lesion Analysis Based Probabilistic Diffusion Tractography

Probabilistic fiber tracking was performed from regions significantly more frequently affected in LG^+^ as compared to LG^-^to localize regions contributing to the deficit in view of intact white matter fiber tracts in elderly subjects. These volumes of interest (VOIs) (Figure 5A) and were affinely transformed from MNI to each participant’s diffusion space (FMRIB’s Linear Image Registration Tool, FLIRT, 12 degrees of freedom; (Jenkinson, Bannister, Brady, & Smith, 2002; Jenkinson & Smith, 2001)). Crossing fiber probabilistic tractography (FDT ProbTrackX) with analysis parameters (step length = 0.5 mm, number of steps = 2000, number of pathways = 5000, curvature threshold = 0.2) previously used to study cerebro-cerebellar-cortical tracts (Salmi et al., 2010), was performed based on each participant’s probability distributions of voxel-wise principal diffusion directions (FDT BedPostX). This algorithm (Behrens, Berg, Jbabdi, Rushworth, & Woolrich, 2007; Behrens et al., 2003) computes the sum of connectivity distributions by generating streamlines from each functionally informed seed mask passing through the respective other mask but excluding pathways that cross into the right hemisphere (sagittal midline exclusion mask at the level of the corpus callosum). Individual patterns of structural connectivity were subdivided into separate bundles by manually placing several inclusion (waypoint) masks to subsequently classify anatomically defined white matter tracts connecting the identified brain regions to cortical and subcortical areas. These left hemisphere inclusion masks were based on the unrestricted overall connectivity pattern of all subjects (crossing fiber probabilistic tractography from both seeds only including a midline exclusion mask) and the ICBM DTI-81 white matter label atlas (http://www.loni.usc.edu; (Mori et al., 2008)). Masks were placed coronally at the level of the left and right middle (cortico-ponto-cerebellar tract) and superior cerebellar peduncle (cerebello-rubro-thalamic tract; (Granziera et al., 2009; Jissendi et al., 2008)), in the left periventricular white matter lateral to the superior corona radiata (superior longitudinal fascicle, SLF; (Makris et al., 2005)), in the left anterior floor of the external capsule (inferior fronto-occipital fasciculus, IFOF; (Catani, Howard, Pajevic, & Jones, 2002)) and in the left posterior corona radiata above the roof of the lateral ventricle (dorsal subcomponent of the IFOF or middle longitudinal fasciculus, MLF; (Makris, Preti, Wassermann, et al., 2013; Martino, Brogna, Robles, Vergani, & Duffau, 2010)). All masks were reverse normalized from MNI standard to individual diffusion space (FLIRT, 12 degrees of freedom). Correct locations of seeds, exclusion and inclusion masks were confirmed visually in native space. All estimated connectivity distributions were scaled across subjects by dividing individual white matter tracts by the total number of probabilistic streamlines to account for differences between tracts due to differences in normalized seed voxel sizes and masks. The tracts were then thresholded to include only voxels that received at least 1 x 10^−7^ percent of the scaled total number of streamlines sent out from the seed masks (samples per voxel (5000) multiplied by the number of voxels in the seed masks (mean = 827.2 ± 61.2 (SD)) (Rilling et al., 2008). Thresholded individual tractography results were binarized, transformed into standard MNI space and averaged to display group variability maps, to quantify the overlap in tract topography. These maps indicate the degree of spatial variability and overlap in each voxel. The pathways and terminations identified were compared against anatomical pathways as defined in primate and human brain dissections or MRI-based anatomical atlases of cortical and subcortical grey or white matter (Oxford thalamic connectivity atlas (Behrens et al., 2003), MNI Talairach atlas (Lancaster et al., 2007; Lancaster et al., 2000), probabilistic cerebellar atlas (Diedrichsen, Balsters, Flavell, Cussans, & Ramnani, 2009), Harvard-Oxford cortical and subcortical probability maps available from FSL (Jenkinson, Beckmann, Behrens, Woolrich, & Smith, 2012)).

## Authors Contributions

A.A. supported tractography data analysis

A.S. conceptualized study, performed experiments, analyzed data, wrote manuscript

D.P. wrote manuscript

M.S. conceptualized study and wrote manuscript

S.A.K. conceptualized study, interpreted data, wrote manuscript

## Acknowledgements

This work was supported by DFG KO 2268/6-1 granted to S. A. K.; A.S. was supported by a dissertation award provided by the University of Leipzig, Germany.

## Notes

### Competing Interest Statement

The authors have declared no competing interest.

## References

**Primary Sources**

Ackermann, H., Graber, S., Hertrich, I., & Daum, I. (1997). Categorical speech perception in cerebellar disorders. Brain and language, 60(2), 323–331. doi:10.1006/brln.1997.1826

Andersen, S. M., Rapcsak, S. Z., & Beeson, P. M. (2010). Cost function masking during normalization of brains with focal lesions: still a necessity? NeuroImage, 53(1), 78–84. doi:10.1016/j.neuroimage.2010.06.003

Ashburner, J., & Friston, K. J. (2005). Unified segmentation. NeuroImage, 26(3), 839–851. doi:10.1016/j.neuroimage.2005.02.018

Aso, K., Hanakawa, T., Aso, T., & Fukuyama, H. (2010). Cerebro-cerebellar interactions underlying temporal information processing. J. Cogn. Neurosci., 22(12), 2913–2925. doi:10.1162/jocn.2010.21429

Baumann, O., Borra, R. J., Bower, J. M., Cullen, K. E., Habas, C., Ivry, R. B., … Sokolov, A. A. (2015). Consensus paper: the role of the cerebellum in perceptual processes. Cerebellum, 14(2), 197–220. doi:10.1007/s12311-014-0627-7

Baumann, O., & Greenlee, M. W. (2007). Neural correlates of coherent audiovisual motion perception. Cerebral cortex, 17(6), 1433–1443. doi:10.1093/cercor/bhl055

Behrens, T. E., Berg, H. J., Jbabdi, S., Rushworth, M. F., & Woolrich, M. W. (2007). Probabilistic diffusion tractography with multiple fibre orientations: What can we gain? NeuroImage, 34(1), 144–155. doi:10.1016/j.neuroimage.2006.09.018

Behrens, T. E., Woolrich, M. W., Jenkinson, M., Johansen-Berg, H., Nunes, R. G., Clare, S., … Smith, S.M. (2003). Characterization and propagation of uncertainty in diffusion-weighted MR imaging. Magnetic resonance in medicine, 50(5), 1077–1088. doi:10.1002/mrm.10609

Berger, K., Weltermann, B., Kolominsky-Rabas, P., Meves, S., Heuschmann, P., Bohner, J., … Buttner, T. (1999). The reliability of stroke scales. The german version of NIHSS, ESS and Rankin scales. Fortschr. Neurol. Psychiatr., 67(2), 81–93. doi:10.1055/s-2007-993985

Bernard, J. A., Peltier, S. J., Benson, B. L., Wiggins, J. L., Jaeggi, S. M., Buschkuehl, M., … Seidler, R. D. (2014). Dissociable functional networks of the human dentate nucleus. Cereb. Cortex, 24(8), 2151–2159. doi:10.1093/cercor/bht065

Boemio, A., Fromm, S., Braun, A., & Poeppel, D. (2005). Hierarchical and asymmetric temporal sensitivity in human auditory cortices. Nat. Neurosci., 8(3), 389–395. doi:10.1038/nn1409

Boersma, P., & Weenink, D. (2001). Praat, a system for doing phonetics by computer. Glot International, 5, 341–345.

Bohsali, A.A., Triplett, W., Sudhyadhom, A., Gullett, J.M., McGregor, K., FitzGerald, D.B., Mareci, T., White, K., & Crosson, B. (2015). Broca’s area - thalamic connectivity. Brain Lang, 141, 80–8. doi: 10.1016/j.bandl.2014.12.001.

Brodal, P. (1978a). The corticopontine projection in the rhesus monkey. Origin and principles of organization. Brain: a journal of neurology, 101(2), 251–283.

Brodal, P. (1978b). Principles of organization of the monkey corticopontine projection. Brain research, 148(1), 214–218.

Brodal, P. (1979). The pontocerebellar projection in the rhesus monkey: an experimental study with retrograde axonal transport of horseradish peroxidase. Neuroscience, 4(2), 193–208.

Callan, D. E., Kawato, M., Parsons, L., & Turner, R. (2007). Speech and song: the role of the cerebellum. Cerebellum, 6(4), 321–327. doi:10.1080/14734220601187733

Catani, M., Howard, R. J., Pajevic, S., & Jones, D. K. (2002). Virtual in vivo interactive dissection of white matter fasciculi in the human brain. NeuroImage, 17(1), 77–94.

Chedru, F., Bastard, V., & Efron, R. (1978). Auditory micropattern discrimination in brain damaged subjects. Neuropsychologia, 16(2), 141–149.

Crinion, J., Ashburner, J., Leff, A., Brett, M., Price, C., & Friston, K. (2007). Spatial normalization of lesioned brains: Performance evaluation and impact on fMRI analyses. NeuroImage, 37(3), 866–875.

De Bleser, R., Cholewa, J., Tabatabaie, N., & Stadie, S. (1997). LeMo, an Expert System for Single Case Assessment of W ord Processing Impairments in Aphasic Patients. Neuropsychological Rehabilitation, 7(4), 339–366. doi:10.1080/713755540

De Bleser, R., Stadie, S., Cholewa, J., & Tabatabaie, N. (2004). LeMo-Lexikon: modellorientierte Einzelfalldiagnostik bei Aphasie, Dyslexie und Dysgraphie. Amsterdam: Elsevier.

Demiralp, T., Basar-Eroglu, C., & Basar, E. (1996). Distributed gamma band responses in the brain studied in cortex, reticular formation, hippocampus and cerebellum. The International journal of neuroscience, 84(1-4), 1–13.

Diedrichsen, J., Balsters, J. H., Flavell, J., Cussans, E., & Ramnani, N. (2009). A probabilistic MR atlas of the human cerebellum. NeuroImage, 46(1), 39–46. doi:10.1016/j.neuroimage.2009.01.045

Dum, R. P., & Strick, P. L. (2003). An unfolded map of the cerebellar dentate nucleus and its projections to the cerebral cortex. Journal of neurophysiology, 89(1), 634–639. doi:10.1152/jn.00626.2002

Efron, R. (1963). Temporal Perception, Aphasia and Deja Vu. Brain: a journal of neurology, 86, 403–424.

Efron, R. (1973). Conservation of temporal information by perceptual systems. Perception & psychophysics, 14(3), 518–530.

Eickhoff, S. B., Stephan, K. E., Mohlberg, H., Grefkes, C., Fink, G. R., Amunts, K., & Zilles, K. (2005). A new SPM toolbox for combining probabilistic cytoarchitectonic maps and functional imaging data. NeuroImage, 25(4), 1325–1335. doi:10.1016/j.neuroimage.2004.12.034

Elangovan, S., & Stuart, A. (2008). Natural boundaries in gap detection are related to categorical perception of stop consonants. Ear and hearing, 29(5), 761–774. doi:10.1097/AUD.0b013e318185ddd2

Fink, M., Churan, J., & Wittmann, M. (2005). Assessment of auditory temporal-order thresholds - a comparison of different measurement procedures and the influences of age and gender. Restorative neurology and neuroscience, 23(5-6), 281–296.

Fink, M., Churan, J., & Wittmann, M. (2006). Temporal processing and context dependency of phoneme discrimination in patients with aphasia. Brain and language, 98(1), 1–11. doi:10.1016/j.bandl.2005.12.005

Flinker, A., Doyle, W. K., Mehta, A. D., Devinsky, O., & Poeppel, D. (2019). Spectrotemporal modulation provides a unifying framework for auditory cortical asymmetries. Nat Hum Behav, 3(4), 393–405. doi:10.1038/s41562-019-0548-z

Frey, S., Campbell, J. S., Pike, G. B., & Petrides, M. (2008). Dissociating the human language pathways with high angular resolution diffusion fiber tractography. The Journal of neuroscience: the official journal of the Society for Neuroscience, 28(45), 11435–11444. doi:10.1523/JNEUROSCI.2388-08.2008

Friederici, A. D. (2011). The brain basis of language processing: from structure to function. Physiological reviews, 91(4), 1357–1392. doi:10.1152/physrev.00006.2011

Granziera, C., Schmahmann, J. D., Hadjikhani, N., Meyer, H., Meuli, R., Wedeen, V., & Krueger, G. (2009). Diffusion spectrum imaging shows the structural basis of functional cerebellar circuits in the human cerebellum in vivo. PloS one, 4(4), e5101. doi:10.1371/journal.pone.0005101

Habas, C., & Cabanis, E. A. (2006). Cortical projections to the human red nucleus: a diffusion tensor tractography study with a 1.5-T MRI machine. Neuroradiology, 48(10), 755–762. doi:10.1007/s00234-006-0117-9

Habas, C., & Cabanis, E. A. (2007). Anatomical parcellation of the brainstem and cerebellar white matter: a preliminary probabilistic tractography study at 3 T. Neuroradiology, 49(10), 849–863. doi:10.1007/s00234-007-0267-4

Hirsh, I. J., & Sherrick, C. E., Jr. (1961). Perceived order in different sense modalities. Journal of experimental psychology, 62, 423–432.

Huber, W., Poeck, K., & Willmes, K. (1984). The Aachen Aphasia Test. Advances in neurology, 42, 291–303.

Ito, M. (2008). Control of mental activities by internal models in the cerebellum. Nature reviews. Neuroscience, 9(4), 304–313. doi:10.1038/nrn2332

Ivry, R. B., & Keele, S. W. (1989). Timing functions of the cerebellum. Journal of cognitive neuroscience, 1(2), 136–152. doi:10.1162/jocn.1989.1.2.136

Ivry, R. B., Spencer, R. M., Zelaznik, H. N., & Diedrichsen, J. (2002). The cerebellum and event timing. Annals of the New York Academy of Sciences, 978, 302–317.

Jenkinson, M., Bannister, P., Brady, M., & Smith, S. (2002). Improved optimization for the robust and accurate linear registration and motion correction of brain images. NeuroImage, 17(2), 825–841.

Jenkinson, M., Beckmann, C. F., Behrens, T. E., Woolrich, M. W., & Smith, S. M. (2012). Fsl. NeuroImage, 62(2), 782–790. doi:10.1016/j.neuroimage.2011.09.015

Jenkinson, M., & Smith, S. (2001). A global optimisation method for robust affine registration of brain images. Medical image analysis, 5(2), 143–156.

Jissendi, P., Baudry, S., & Baleriaux, D. (2008). Diffusion tensor imaging (DTI) and tractography of the cerebellar projections to prefrontal and posterior parietal cortices: a study at 3T. Journal of neuroradiology. Journal de neuroradiologie, 35(1), 42–50. doi:10.1016/j.neurad.2007.11.001

Keele, S. W., & Ivry, R. (1990). Does the cerebellum provide a common computation for diverse tasks? A timing hypothesis. Annals of the New York Academy of Sciences, 608, 179–207.

Kelly, R. M., & Strick, P. L. (2003). Cerebellar loops with motor cortex and prefrontal cortex of a nonhuman primate. The Journal of neuroscience: the official journal of the Society for Neuroscience, 23(23), 8432–8444.

Kotz, S. A., & Schwartze, M. (2010). Cortical speech processing unplugged: a timely subcortico-cortical framework. Trends in cognitive sciences, 14(9), 392–399. doi:10.1016/j.tics.2010.06.005

Kotz, S. A., Stockert, A., & Schwartze, M. (2014). Cerebellum, temporal predictability and the updating of a mental model. Philosophical transactions of the Royal Society of London. Series B, Biological sciences, 369(1658), 20130403. doi:10.1098/rstb.2013.0403

Lancaster, J. L., Tordesillas-Gutierrez, D., Martinez, M., Salinas, F., Evans, A., Zilles, K., … Fox, P. T. (2007). Bias between MNI and Talairach coordinates analyzed using the ICBM-152 brain template. Human brain mapping, 28(11), 1194–1205. doi:10.1002/hbm.20345

Lancaster, J. L., Woldorff, M. G., Parsons, L. M., Liotti, M., Freitas, C. S., Rainey, L., … Fox, P. T. (2000). Automated Talairach atlas labels for functional brain mapping. Human brain mapping, 10(3), 120–131.

Levitt, H. (1971). Transformed up-down methods in psychoacoustics. The Journal of the Acoustical Society of America, 49(2), 467–477.

Liebermann, D., Ploner, C. J., Kraft, A., Kopp, U. A., & Ostendorf, F. (2013). A dysexecutive syndrome of the medial thalamus. Cortex; a journal devoted to the study of the nervous system and behavior, 49(1), 40–49. doi:10.1016/j.cortex.2011.11.005

Makris, N., Kennedy, D. N., McInerney, S., Sorensen, A. G., Wang, R., Caviness, V. S., Jr., & Pandya, D. N. (2005). Segmentation of subcomponents within the superior longitudinal fascicle in humans: a quantitative, in vivo, DT-MRI study. Cerebral cortex, 15(6), 854–869. doi:10.1093/cercor/bhh186

Makris, N., Preti, M. G., Asami, T., Pelavin, P., Campbell, B., Papadimitriou, G. M., … Kubicki, M. (2013). Human middle longitudinal fascicle: variations in patterns of anatomical connections. Brain structure & function, 218(4), 951–968. doi:10.1007/s00429-012-0441-2

Makris, N., Preti, M. G., Wassermann, D., Rathi, Y., Papadimitriou, G. M., Yergatian, C., … Kubicki, M. (2013). Human middle longitudinal fascicle: segregation and behavioral-clinical implications of two distinct fiber connections linking temporal pole and superior temporal gyrus with the angular gyrus or superior parietal lobule using multi-tensor tractography. Brain imaging and behavior, 7(3), 335–352. doi:10.1007/s11682-013-9235-2

Martino, J., Brogna, C., Robles, S. G., Vergani, F., & Duffau, H. (2010). Anatomic dissection of the inferior fronto-occipital fasciculus revisited in the lights of brain stimulation data. Cortex; a journal devoted to the study of the nervous system and behavior, 46(5), 691–699. doi:10.1016/j.cortex.2009.07.015

Molholm, S., Sehatpour, P., Mehta, A. D., Shpaner, M., Gomez-Ramirez, M., Ortigue, S., … Foxe, J. J. (2006). Audio-visual multisensory integration in superior parietal lobule revealed by human intracranial recordings. Journal of neurophysiology, 96(2), 721–729. doi:10.1152/jn.00285.2006

Mori, S., Oishi, K., Jiang, H., Jiang, L., Li, X., Akhter, K., Mazziotta, J. (2008). Stereotaxic white matter atlas based on diffusion tensor imaging in an ICBM template. NeuroImage, 40(2), 570–582. doi:10.1016/j.neuroimage.2007.12.035

Oldfield, R. C. (1971). The assessment and analysis of handedness: the Edinburgh inventory. Neuropsychologia, 9(1), 97–113.

Pal, D., Trivedi, R., Saksena, S., Yadav, A., Kumar, M., Pandey, C. M., … Gupta, R. K. (2011). Quantification of age- and gender-related changes in diffusion tensor imaging indices in deep grey matter of the normal human brain. Journal of clinical neuroscience: official journal of the Neurosurgical Society of Australasia, 18(2), 193–196. doi:10.1016/j.jocn.2010.05.033

Pandya, D. N., Rosene, D. L., & Doolittle, A. M. (1994). Corticothalamic connections of auditory-related areas of the temporal lobe in the rhesus monkey. The Journal of comparative neurology, 345(3), 447–471. doi:10.1002/cne.903450311

Pastor, M. A., Artieda, J., Arbizu, J., Marti-Climent, J. M., Penuelas, I., & Masdeu, J. C. (2002). Activation of human cerebral and cerebellar cortex by auditory stimulation at 40 Hz. The Journal of neuroscience: the official journal of the Society for Neuroscience, 22(23), 10501–10506.

Pastor, M. A., Macaluso, E., Day, B. L., & Frackowiak, R. S. (2006). The neural basis of temporal auditory discrimination. NeuroImage, 30(2), 512–520. doi:10.1016/j.neuroimage.2005.09.053

Pastor, M. A., Thut, G., & Pascual-Leone, A. (2006). Modulation of steady-state auditory evoked potentials by cerebellar rTMS. Experimental Brain Research, 175(4), 702–709. doi:10.1007/s00221-006-0588-2

Pastor, M. A., Vidaurre, C., Fernandez-Seara, M. A., Villanueva, A., & Friston, K. J. (2008). Frequency-specific coupling in the cortico-cerebellar auditory system. Journal of neurophysiology, 100(4), 1699–1705. doi:10.1152/jn.01156.2007

Petacchi, A., Laird, A. R., Fox, P. T., & Bower, J. M. (2005). Cerebellum and auditory function: an ALE meta-analysis of functional neuroimaging studies. Human brain mapping, 25(1), 118–128. doi:10.1002/hbm.20137

Petersen, K.J., Reid, J.A., Chakravorti, S., Juttukonda, M.R., Franco, G., Trujillo, P., Stark, A.J., Dawant, B.M., Donahue, M.J., & Claassen, D.O. (2018). Structural and functional connectivity of the nondecussating dentato-rubro-thalamic tract. Neuroimage, 176, 364–371. doi: 10.1016/j.neuroimage.2018.04.074.

Poeppel, D. (2003). The analysis of speech in different temporal integration windows: cerebral lateralization as ‘asymmetric sampling in time’. Speech Communication, 41(1), 245–255. doi: 10.1016/S0167-6393(02)00107-3

Pöppel, E. (1978). Time Perception. In R. Held, H. W. Leibowitz, & H.-L. Teuber (Eds.), Perception (pp. 713–729). Berlin, Heidelberg, Germany: Springer Berlin Heidelberg.

Powell, J. L., Parkes, L., Kemp, G. J., Sluming, V., Barrick, T. R., & Garcia-Finana, M. (2012). The effect of sex and handedness on white matter anisotropy: a diffusion tensor magnetic resonance imaging study. Neuroscience, 207, 227–242. doi:10.1016/j.neuroscience.2012.01.016

Ramnani, N. (2006). The primate cortico-cerebellar system: anatomy and function. Nature reviews. Neuroscience, 7(7), 511–522. doi:10.1038/nrn1953

Ramnani, N., Behrens, T. E., Johansen-Berg, H., Richter, M. C., Pinsk, M. A., Andersson, J. L., … Matthews, P. M. (2006). The evolution of prefrontal inputs to the cortico-pontine system: diffusion imaging evidence from Macaque monkeys and humans. Cerebral cortex, 16(6), 811–818. doi:10.1093/cercor/bhj024

Rilling, J. K., Glasser, M. F., Preuss, T. M., Ma, X., Zhao, T., Hu, X., & Behrens, T. E. (2008). The evolution of the arcuate fasciculus revealed with comparative DTI. Nature neuroscience, 11(4), 426–428. doi:10.1038/nn2072

Robson, H., Grube, M., Lambon Ralph, M. A., Griffiths, T. D., & Sage, K. (2013). Fundamental deficits of auditory perception in Wernicke’s aphasia. Cortex; a journal devoted to the study of the nervous system and behavior, 49(7), 1808–1822. doi:10.1016/j.cortex.2012.11.012

Rorden, C., Bonilha, L., Fridriksson, J., Bender, B., & Karnath, H. O. (2012). Age-specific CT and MRI templates for spatial normalization. NeuroImage, 61(4), 957–965. doi:10.1016/j.neuroimage.2012.03.020

Rorden, C., & Karnath, H. O. (2004). Using human brain lesions to infer function: a relic from a past era in the fMRI age? Nature reviews. Neuroscience, 5(10), 813–819. doi:10.1038/nrn1521

Rorden, C., Karnath, H. O., & Bonilha, L. (2007). Improving lesion-symptom mapping. Journal of cognitive neuroscience, 19(7), 1081–1088. doi:10.1162/jocn.2007.19.7.1081

Rosen, S. (1992). Temporal information in speech: acoustic, auditory and linguistic aspects. Philosophical transactions of the Royal Society of London. Series B, Biological sciences, 336(1278), 367–373. doi:10.1098/rstb.1992.0070

Salat, D. H., Tuch, D. S., Greve, D. N., van der Kouwe, A. J., Hevelone, N. D., Zaleta, A. K., … Dale, A. M. (2005). Age-related alterations in white matter microstructure measured by diffusion tensor imaging. Neurobiology of aging, 26(8), 1215–1227. doi:10.1016/j.neurobiolaging.2004.09.017

Salmi, J., Pallesen, K. J., Neuvonen, T., Brattico, E., Korvenoja, A., Salonen, O., & Carlson, S. (2010). Cognitive and motor loops of the human cerebro-cerebellar system. Journal of cognitive neuroscience, 22(11), 2663–2676. doi:10.1162/jocn.2009.21382

Saur, D., Kreher, B. W., Schnell, S., Kummerer, D., Kellmeyer, P., Vry, M. S., … Weiller, C. (2008). Ventral and dorsal pathways for language. Proceedings of the National Academy of Sciences of the United States of America, 105(46), 18035–18040. doi:10.1073/pnas.0805234105

Schmahmann, J., & Pandya, D. N. (2009). Superior temporal region. In J. Schmahmann & D. N. Pandya (Eds.), Fiber Pathways of the Brain (pp. 143–186). New York, New York: Oxford University Press.

Schmahmann, J. D. (2004). Disorders of the cerebellum: ataxia, dysmetria of thought, and the cerebellar cognitive affective syndrome. The Journal of neuropsychiatry and clinical neurosciences, 16(3), 367–378. doi:10.1176/appi.neuropsych.16.3.367

Schmahmann, J. D., & Pandya, D. N. (1991). Projections to the basis pontis from the superior temporal sulcus and superior temporal region in the rhesus monkey. The Journal of comparative neurology, 308(2), 224–248. doi:10.1002/cne.903080209

Schwartze, M., & Kotz, S. A. (2013). A dual-pathway neural architecture for specific temporal prediction. Neuroscience and biobehavioral reviews, 37(10 Pt 2), 2587–2596. doi:10.1016/j.neubiorev.2013.08.005

Schwartze, M., & Kotz, S. A. (2016). Contributions of cerebellar event-based temporal processing and preparatory function to speech perception. Brain and language, 161, 28–32. doi:10.1016/j.bandl.2015.08.005

Schwartze, M., Tavano, A., Schröger, E., & Kotz, S. A. (2012). Temporal aspects of prediction in audition: cortical and subcortical neural mechanisms. International journal of psychophysiology: official journal of the International Organization of Psychophysiology, 83(2), 200–207. doi:10.1016/j.ijpsycho.2011.11.003

Sokolov, A. A., Erb, M., Grodd, W., & Pavlova, M. A. (2014). Structural loop between the cerebellum and the superior temporal sulcus: evidence from diffusion tensor imaging. Cerebral cortex, 24(3), 626–632. doi:10.1093/cercor/bhs346

Spencer, R. M. C., & Ivry, R. B. (2013). Cerebellum and Timing. In M. Manto, J. D. Schmahmann, F. Rossi, D. L. Gruol, & N. Koibuchi (Eds.), Handbook of the cerebellum and cerebellar disorders (pp. 1201–1219). New York: Springer.

Swisher, L., & Hirsh, I. J. (1972). Brain damage and the ordering of two temporally successive stimuli. Neuropsychologia, 10(2), 137–152.

Turken, A. U., & Dronkers, N. F. (2011). The neural architecture of the language comprehension network: converging evidence from lesion and connectivity analyses. Frontiers in systems neuroscience, 5, 1. doi:10.3389/fnsys.2011.00001

Wang, Y., Fernandez-Miranda, J. C., Verstynen, T., Pathak, S., Schneider, W., & Yeh, F. C. (2013). Rethinking the role of the middle longitudinal fascicle in language and auditory pathways. Cerebral cortex, 23(10), 2347–2356. doi:10.1093/cercor/bhs225

Weise, A., Bendixen, A., Muller, D., & Schröger, E. (2012). Which kind of transition is important for sound representation? An event-related potential study. Brain research, 1464, 30–42. doi:10.1016/j.brainres.2012.04.046

Wolpert, D. M., Ghahramani, Z., & Jordan, M. I. (1995). An internal model for sensorimotor integration. Science, 269(5232), 1880–1882.

Wolpert, D. M., & Miall, R. C. (1996). Forward Models for Physiological Motor Control. Neural networks: the official journal of the International Neural Network Society, 9(8), 1265–1279.

Wright, H. N. (1960). Audibility of Switching Transients. Journal of the Acoustical Society of America, 32, 138. doi:10.1121/1.1907866

Yund, E. W., & Efron, R. (1974). Dichoptic and dichotic micropattern discrimination. Perception & psychophysics, 15(2), 383–390.

Zatorre, R. J., & Belin, P. (2001). Spectral and temporal processing in human auditory cortex. Cerebral cortex, 11(10), 946–953.

